# Tracking Cell Layer Contribution During Repair of the Tympanic Membrane

**DOI:** 10.1101/2023.08.09.552665

**Authors:** Olivia M. Dinwoodie, Abigail S. Tucker, Juan Fons-Romero

**Affiliations:** Centre for Craniofacial and Regenerative Biology, Faculty of Dentistry, Oral and Craniofacial Sciences, King’s College London, London SE1 9RT, UK

**Keywords:** ear drum, proliferation, wound healing, germ layer

## Abstract

The tympanic membrane (or ear drum) is found at the interface between the middle ear and the external ear. The membrane is composed of three layers of different embryonic origin: an outer ectodermally-derived layer, a middle neural crest-derived fibroblast layer with contribution from the mesoderm-derived vasculature, and an inner endodermally-derived mucosal layer. These layers form a thin sandwich which is often perforated as a consequence of trauma, pressure changes, or middle ear inflammation. Usually, the tympanic membrane heals with minimal scarring, but in 6% of cases the perforation fails to heal leading to hearing loss, tinnitus and pain requiring surgery. How cells bridge the gap to close the perforation is an interesting question, as this needs to happen in the absence of an initial scaffold. Here we assess the contribution, timing, and interaction of the different layers of the membrane during repair in the mouse using markers and reporter mouse lines. We show that the ectodermal layer retracts after perforation, before proliferating away from the wound edge, with Keratin 5 basal cells migrating over the hole to bridge the gap. The mesenchymal and mucosal layers then use this scaffold to complete the repair, in tandem with changes in the vasculature. Finally, differentiation of the epithelium leads to formation of a scab that falls off. Our results reveal the dynamics and interconnections between the embryonic germ layers during repair and highlight how defects in healing may occur. Unearthing the complexities of TM healing is important as chronic TMP is a common clinical issue with limited treatment options.

## Introduction

The tympanic membrane (TM) is located between the external and the middle ear, where sound vibrations are picked up and transmitted via the manubrial insertion at the center of the TM through the ossicular chain and to the cochlea. The TM has two main parts, the pars tensa (PT) which is the site of sound conduction, and the pars flaccida (PF) the role of which is thought to involve pressure equilibration (Lim, 1995; Robert et al., 1982) (Figure 1A). The PT houses the annulus; the thicker outer ring of the TM, as well as the manubrium of the malleus, which is where the TM connects to the ossicular chain of the middle ear. The PT is stretched taut, held by the tympanic ring, with the outer and inner layers continuous with the external ear canal and mucosa of the middle ear respectively (Tucker et al., 2018).

**Fig.1.**
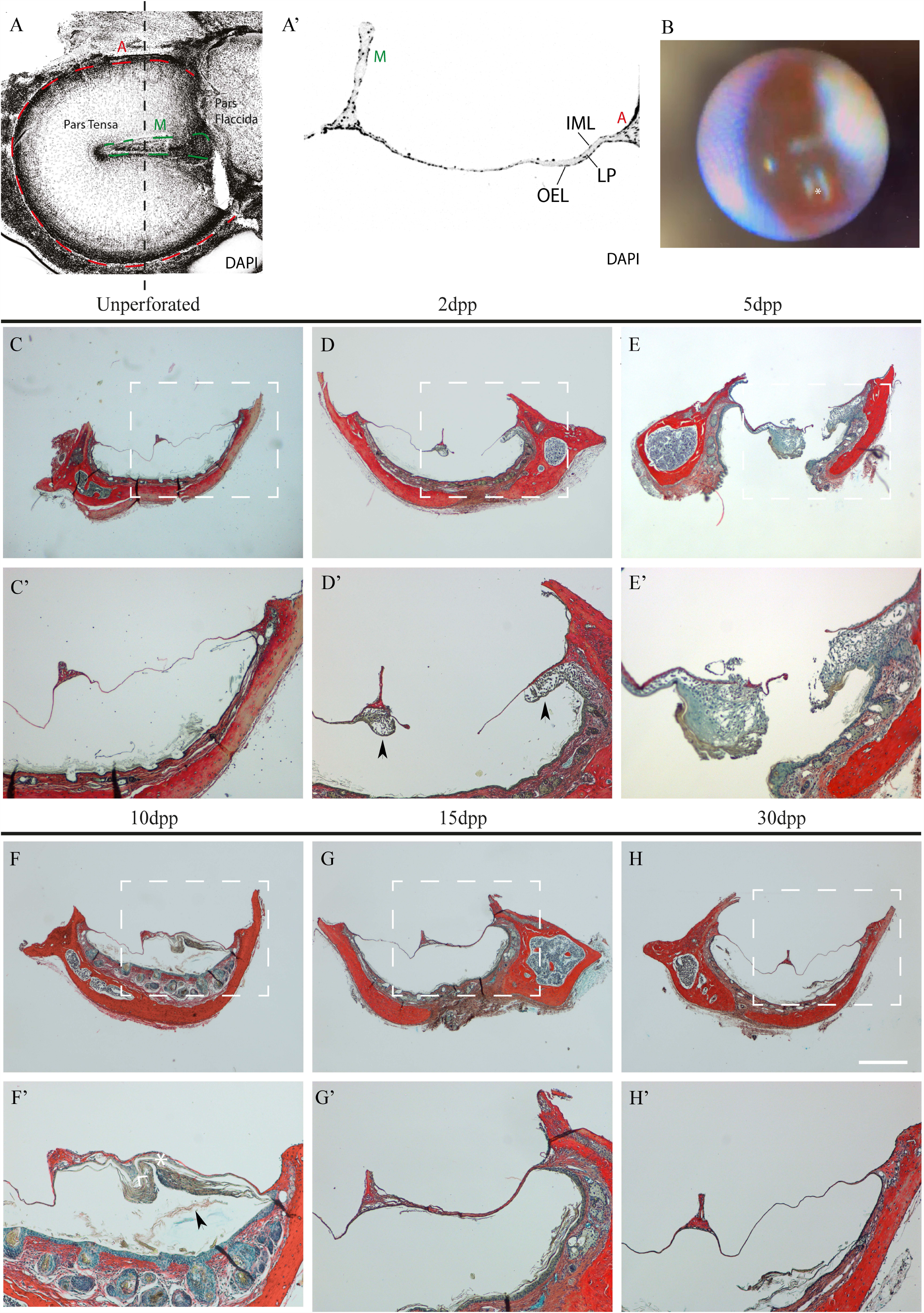
The tympanic membrane heals by 30 days post-perforation. (A) whole mount view of a dissected tympanic membrane imaged from the middle ear side, showing the Pars Tensa, Pars Flaccida, manubrium (M) and annulus (A). Nuclei (DAPI) is showed in grey. (A’) Transversal section at the level indicated in A by the dotted line. It is showed the manubrium (M), the inner mucosal layer (IML), the lamina propria (LP), the outer epithelial layer (OEL) and the annulus (A). (B) Otoscopy photograph of a perforated tympanic membrane (asterisk marks the site of perforation). (C-H) Trichrome staining of the timeline of healing. (C,C’) Unperforated tympanic membrane. (D,D’) 2 days post-perforation (2dpp). Arrowheads highlight regions of tissue expansion away from the wound edge. (E,E’) 5dpp (F,F’),10dpp (G,G’) 15dpp (H,H) 30dpp. Scale Bar: H = 500μm (same scale A-G). Scale bar H’ = 100μm (same scale A’-G’).

The TM represents a unique example of a single tissue derived from all three germ layers (Takechi et al., 2016). The outer epidermal layer, derived from the ectoderm, consists of keratinocytes. These vary according to their extent of differentiation and resemble a cellular hierarchy reminiscent of the cell types in the epidermis of the skin (Frumm et al., 2021). The underlying lamina propria consists of neural crest-derived mesenchyme with contribution from the mesoderm-derived blood vessels and nerves of the TM. The neural-crest derived mesenchyme and can be identified via *Wnt1* lineage tracing (Chai et al., 2000). Finally, the inner mucosal layer is endodermally-derived and can be identified by *Sox17* lineage tracing (Thompson & Tucker, 2013; Nowotschin et al., 2019). The lamina propria is thickened towards the annulus and manubrium and extremely thin towards the centre of the PT (Figure 1A’). The PT and PF have a similar layered structure but differ in the make-up of their lamina proprias. In the PT, this layer contains highly assembled radial and circular collagen fibres, consisting mainly of type II collagen, whereas type I collagen is found more loosely arranged in the PF (Stenfeldt et al., 2006, Knutsson et al., 2007).

During development, ectodermal cells from the region of the first pharyngeal arch invaginate towards the endodermal out-pocketing of the first pharyngeal pouch (Minoux et al., 2013). The bones of the middle ear form from the neural crest-derived mesenchyme (Richter et al., 2010). As the ectodermally derived external acoustic meatus invaginates (Fons et al., 2020) and the endodermal middle ear cavity extends, the two layers enclose a layer of mesenchyme between them ultimately resulting in the typical trilaminar structure of the TM (Mallo, 2001).

Acute eardrum perforation is a major clinical presentation and can result from pressure changes, trauma, and infection. In 94% of cases, this perforation can heal itself with no scarring and no changes to original hearing capability. Conversely, 6% of cases do not heal and become chronic, posing a strain on patients and health services worldwide (Dolhi & Weimer, 2022).

Previous studies have revealed a general account of TMP healing in a range of mammalian species, including guineapig, cat, rat, and mouse (Gladstone et al., 1995; Stenfors et al., 1980). Histological analysis in the rat has suggested the potential timing of the reaction from each of the layers, with an initial epithelial response, followed by the middle and inner layers (de Araújo et al., 2014). In the cat, the perforation edge was found to thicken with an epithelial bridge observed 7-9 days after perforation (Stenfors et al., 1980). A retraction of the epithelium was noted 48hrs after perforation, leaving an uncovered middle layer (Stenfors et al., 1980). A similar retraction was observed in histological section in the rat, with collagen fibers left exposed (de Araújo et al., 2014). However, due to the thinness of the membrane, it is difficult to follow the contribution of the different layers without molecular markers.

In intact ear drums, proliferative centers have been noted to form around the annulus and near the manubrium (Chari et al., 2019; Tucker et al., 2018). Following perforation, a large increase in the number of proliferating cells have been shown in all three layers, but highest in the ectoderm layer (Chari et al., 2019). Interestingly, similar to homeostasis, proliferation after perforation was focused around the annulus and the malleus (Chari et al., 2019). Proliferation at a distance from the wound site has also been noted in the guineapig, suggesting that epithelial cells migrate to the wound site from distinct regions (Clawson & Litton, 1971; Reeve, 1977). Ink tracing experiments have suggested that the basal and spinosum layers contribute to repair (Gladstone et al., 1995), with high expression of the basal marker Keratin5, upregulated during repair (Chari et al., 2019;). If proliferation is inhibited using Mitomycin C in rats, the perforations fail to heal even after 2 months of study, highlighting the essential role of proliferation (Guneri et al., 2003).

The phases of TM wound healing have been classified into stages using microarray analysis, mirroring the phases noted during skin wound healing (Santa Maria et al., 2011). Immediately following perforation is the inflammatory phase, whereby genes involved in acute inflammation are upregulated, as well as some growth factors (e.g., EGF) (Santa Maria et al., 2011). This phase takes place almost immediately following perforation to 2dpp (days post perforation). Following the inflammatory phase, the proliferative phase (2-4/5 dpp) initiates with upregulation of signaling pathways, such as FGFs and PDGF (Santa Maria et al., 2011). Finally, a remodeling phase occurs (7-14dpp), whereby thinning of the tissue takes place (Santa Maria et al., 2011). It is worth noting that while TM healing is often compared to skin wound healing there are differences. Importantly, skin wound healing requires an initial scaffold of fibroblasts, creating a platform for re-epithelialization, while the TM appears to involve migration of keratinocytes to bridge the gap and to create an initial scaffold, along which the fibroblasts migrate (Gladstone et al., 1995). Interestingly, one study labelling TM perforations with ink, suggested that the inner mucosal layer formed the initial bridge across the perforation, along which the ectodermal cells then migrated (Johnson & Hawke, 1986). The order in which the layers migrate is therefore unclear. That the middle, fibroblast layer arises last is supported by studies in humans where the perforation is replaced by a bi-layered membrane without the fibroblast layer, or a membrane with disorganized fibrils (Yamashita, 1985).

While the focus of previous work has rested on the presence of potential stem cell populations and surgical options for enhancing repair, further investigation is required to understand the cellular contribution, precise timing, and mechanisms of the healing process. This paper uses transgenic reporter mouse models to show how cellular components from each of the three layers of the TM contribute to the repair of the tissue, uncovering novel parts of the healing process, and highlighting the timing of reaction for each cell layer. We show that the epithelial layer initially retracts post-perforation, leaving behind both the inner and middle layers. The retraction is followed by the formation of the epithelial scaffold, which is composed through localized areas of proliferating K5+ve keratinocytes, which later mature. We show that the mesenchymal and endodermal layers respond later with proliferation and migration in tandem with changes to the vasculature. Understanding the mechanisms of repair in the TM is crucial, as without this knowledge we cannot begin to design treatments to enhance repair in acute as well as chronic perforated TMs.

## Materials and methods

### Animals

All mice were kept in the Biological Services Unit at King’s College London. All animal husbandry and procedures were carried out in accordance with King’s College ethical guidelines and Home Office guidelines. *Wnt1Cre* (Danielian et al., 1998), Mesp1Cre (Saga et al., 1999), *Sox17-2A-iCre* (Engert et al., 2009) males and Rosa26;*Tomato* females (*tdTomato [Gt(ROSA)26 Sor tm14(CAG-tdTomato); Hze JAX labs*]) were bred to create the reporter mouse lines: *Wnt1cre;tomato, Mesp1cre;tomato* and *Sox17icre;tomato*.

### Perforations

Ear drum perforation was performed in 6-8 weeks old wild-type CD1 (N = 20), Wnt1cre;tomato (N = 13), Mesp1cre;tomato (N = 8) and Sox17cre;tomato (N = 5) adult mice. Transgenic mice were on a mixed CD1/C57bl6 background. Both male and female mice were used for experimentation. All ear drums healed by 15 days with no difference shown for sex. Perforations were created using a 27g syringe and 1mm endoscopic camera for visualisation (Figure 1B)(RVA synergies) while animals were under anaesthesia with a cocktail of ketamine and medetomidine at 75mg/kg. After surgery, anaesthesia was reversed using antipedamizole and buprenorphine. At specific time points mice were sacrificed using approved schedule one culling methods, and the ear drums, attached to the tympanic ring, dissected out under a Leica MZFLIII dissection scope. Dissected tissue was fixed in 4% paraformaldehyde for one hour and DAPI stained prior to whole-mount imaging. All animal surgery was approved by the UK Home office with licences in place.

### Histology and Immunohistochemistry

For paraffin embedded sectioning, samples were decalcified in 0.5M EDTA (pH 9), dehydrated via an ethanol series (PBS, 30%, 50%, 70%,90%, 100%), cleared using xylene and embedded in paraffin wax. 6um sections were cut using a Leica microtome RM2245 and mounted onto superfrost slides (FisherScientific 11976299). Immunofluorescence was performed via standard protocols (Zaqout et al., 2020). The paraffin slides were initially dewaxed in xylene (3x 10 mins) and rehydrated via an ethanol series (100%,90%,70%,50%,30%, PBS 2 mins each). Antigen retrieval was achieved using 0.1M citrate buffer (pH 6) in a water bath at 95°C. After antigen retrieval slides were washed with PBS and blocked using blocking buffer (10% FBS, 1% BSA and 0.0125% tween20 in PBS for 1-2 hours). Primary antibodies were then applied overnight at 4°C (see supplementary material for table of antibodies). The primary antibodies were washed using PBT (4x5 mins), before secondary antibodies (including DAPI) made up in blocking solution, were applied. Finally, slides were washed in PBT (4x5 minutes), mounted using fluoroshield (Sigma F6182) and left to dry overnight before imaging. TM sections are fragile, and optimisations of the immunofluorescence protocol included: Not preheating antigen retrieval buffer and using 0.0125% Tween20 in PBT when washing and in blocking solution.

This protocol was followed for all immunofluorescence experiments apart from that for Collagen (II) which requires a pre-antigen retrieval step whereby slides were treated with 3% hydrogen peroxide for 30 mins at RTP. Slides were also processed via a trichrome stain with Sirrius red, Ehrlich’s haematoxylin and Alcian blue under standard protocols. Immunofluorescence experiments were repeated on 2-3 TM samples.

### Imaging

Whole ear drums on FluoroDishes (FD35-100), or immunofluorescent stained slides were imaged on a Leica TCS SP5 confocal microscope with LAS AF software. Trichrome stained slides were imaged on the Nikon Eclipse 80i and brightfield images were taken using the Leica FlexaCam A5. Images were processed and quantified in Image J (version 1.0) and figures (including schematics) were made using Adobe Illustrator (2021) or Adobe Photoshop (2021). Graphs were made using GraphPad Prism 9.

## Results

### Small tympanic membrane perforations resolve over 30 days in the mouse

To understand the timing of the healing process, repair was characterised in wildtype mice using histological analysis from perforation to 30 days post-operation. Perforations were consistently placed in the right lower quadrant of the Pars Tensa (PT) in between the manubrium and annulus, with an endoscopic camera utilised to allow visualization during perforation (Figure 1B). Two days after perforation a clear hole was present, compared to unperforated ears, with a pronounced thickening of the membrane at a distance from the injury near to the annulus and manubrium (Fig. 1C,D, C’,D’). 5 days after injury, the membrane continued to thicken on the outside of the ear drum. At this stage approximately 50% of ears analysed still showed a centre hole where the perforation had been made, while a bridge over the hole was evident in the other 50% (Fig.1E, E’ and data not shown). By 10 days, a bridge had formed over the injury site with evidence of keratinisation and shedding of cells into the external ear canal (Fig.1F,F’). After 15dpp, the membrane was almost back to normal thickness compared to the uninjured side of the membrane (Fig.1G, G’), and by 30dpp there were no overt signs of any previous damage (Fig.1H, H’).

### The epidermal layer retracts leaving behind the middle and inner layers

The histological analysis at 2 days post perforation (2dpp) suggested a retraction of tissue away from the initial wound site. A similar early retraction of tissue away from the perforation was previously observed in the rat and cat (Reeve, 1977; Stenfors et al., 1980), but has not been described in more recent literature. In wholemount, the retraction was very evident at 2 days (Fig. 2B,C). The tympanic membrane is a taut structure, held tightly within the bony tympanic ring, it was therefore hypothesised that the TM retraction may have been a mechanical consequence of this tension and, therefore, occur immediately post-perforation. This was tested by imaging the retracted membrane 3 hours after perforation (Fig. 2A). In contrast to the large retracted area at 2-days post perforation, there was no evidence of retraction at 3hrspp (Fig. 2A). Retraction of tissue away from the wound site was, therefore, not immediate due to the release of mechanical tension within the drum. Perforated TMs were stained with phalloidin, which labels F-actin cables, to outline the cells in the retracted and un-retracted regions at 2dpp (Fig.2C). High phalloidin levels were evident around the retracted tissue but not in the cells left behind around the perforation site (Fig. 2C). The area of the perforation was measured compared to the area retracted (Fig2.D) (N = 8), showing a very consistent area of retraction, approximately 3-4 times the area of the original perforation (Fig.2D).

**Fig.2.**
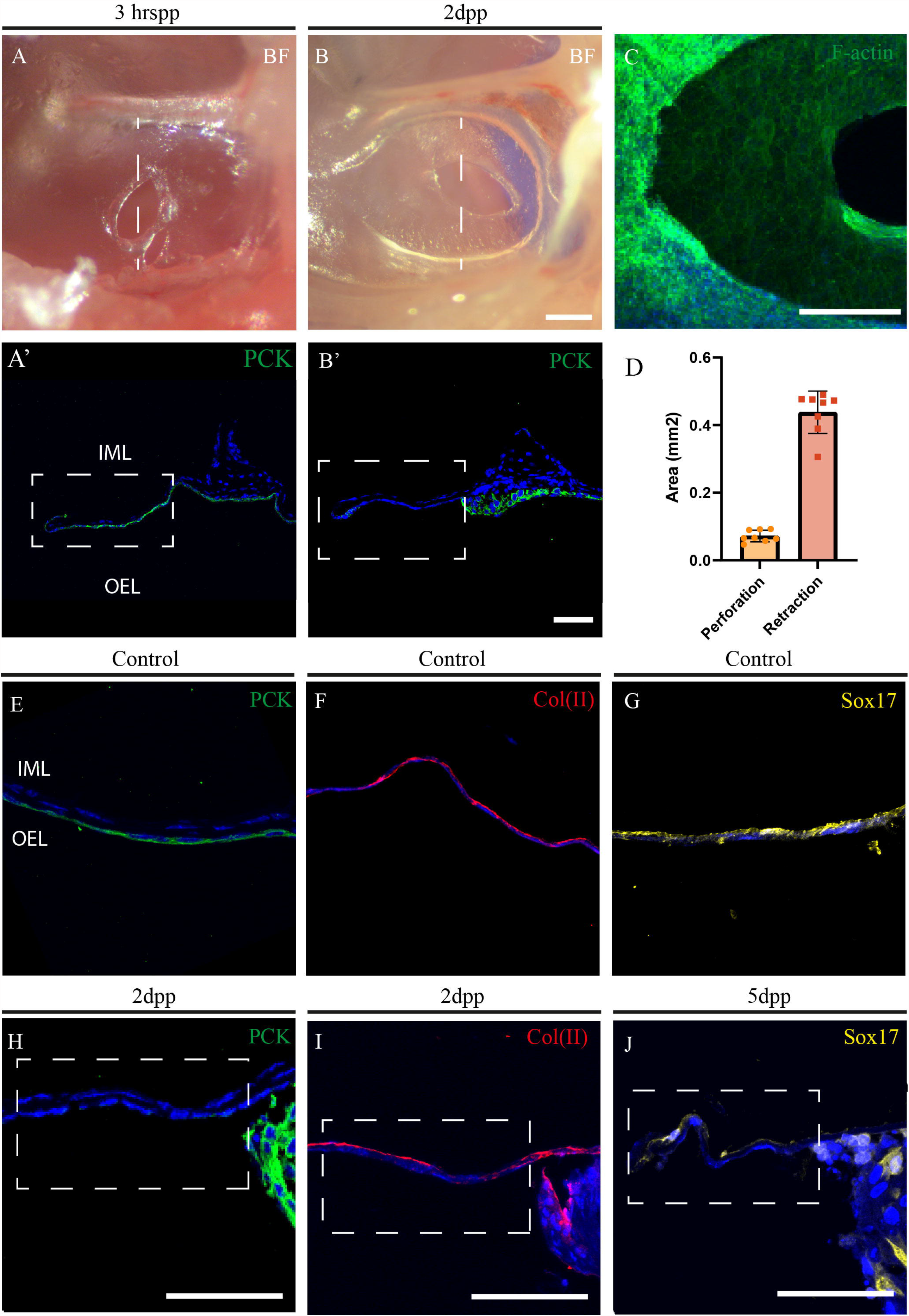
Retraction of the outer epithelial layer post-perforation. A,B)Bright field picture of a tympanic membrane 3hr (A) and 2days (B) after perforation. A’,B’) IF of a transversal section taken at the level of the dotted line in A and B respectively showing the epithelial marker Pancytokeratin in green. C) whole mount fluorescence of actin filaments with phalloidin in green. D) graph of the area measurement of the perforation and the retraction of the outer epithelial layer. E-J) IF for pancytokeratin (E,H), Collagen II (F,I) and Tomato (G,J) in the reporter mouse *Sox17icre-tdTomato*; before (E,F,G) and after (H,I,J) perforation. Dpp: days post perforation. Scale bar: B = 200μm (same scale in A). B’ and H, I,J = 50μm (same scale in A’). Scale bar in C = 200μm.

In the cat and rat retraction has been suggested to be due to pulling away of the outer epithelial layer. To confirm which layer had retracted, TMs were sectioned (Fig.2 A,B dotted lines) and stained for pan-cytokeratin (PCK), a marker of the outer epidermal layer (Fig. 2E). At 3hrspp PCK positive cells were observed in the outer layer reaching the edge of the perforation (Fig2A’), while by 2dpp the domain of PCK expression was limited to the retracted membrane (Fig.2B’,H). The outer ectodermally-derived epithelium, therefore, retracts as part of the early wound response. Having shown the retraction of the ectoderm after perforation, the nature of the tissue layers that remained around the perforation were analysed. Collagen II was used as a marker of the lamina propria (LP) to discern whether the un-retracted membrane was composed of middle layer tissue (Fig. 2F). In contrast to the loss of PCK (Fig.2H), Collagen II staining was observed in the un-retracted membrane (Fig.2I), suggesting that part of the membrane ‘left behind’ was the LP. As the inner mucosal layer is of endodermal embryonic origin, a Sox17icre reporter mouse was used to identify endodermally derived tissues in the adult at 5dpp (Fig. 2G). Positive staining was evident in the un-retracted membrane (Fig.2J), highlighting that while the outer ectoderm layer retracted away from the wound site after injury, the mesenchymal and endodermal layers remained at the wound site.

### Basal epithelial cells cover the wound and create a scaffold

In order to investigate the contribution of the epithelial layer to the healing of the perforated TM, three epithelial markers of specific stages of keratinocyte differentiation were analysed. Keratin 5 (K5) marks undifferentiated basal epithelial stem cells, while Keratin 10 (K10) and Loricrin mark committed and terminally differentiated keratinocytes respectively (Koster & Roop, 2004). The unperforated ear drum has been shown to have a three-dimensional differentiation hierarchy of keratinocytes, with clusters of undifferentiated and differentiated cells (Frumm et al., 2021). In keeping with this, in sections through unperforated TMs, Keratin 5, Keratin 10 and Loricrin positive cells were present within the ectodermal layer (Fig.3A, E, I). Between 5 and 7 d pp, the epithelium had expanded and created a scaffolding covering the perforated site (Fig. 3B). The newly constructed bridge was K5 positive (Fig.3B), but, unlike the ear drum during homeostasis, contained no K10 and Loricrin positive cells (Fig.3F,J). Healing of the wound therefore involves the undifferentiated, basal cell population. By 10dpp a pronounced reduction of K5 positive cells was observed (Fig.3C) as well as the appearance of committed (K10+ve) keratinocytes (Fig.3G) and terminally differentiated (Loricrin+ve) keratinocytes (Fig.3K), indicating that the ells involved in the initial epithelial ingrowth had started to differentiate. The presence of Loricrin positive keratinocytes at day 10-15 marked the site where the membrane would eventually peel off in layers to achieve the thin 1-2 cell layers of the healed membrane (Fig.3 K, L). There was evidence of cornification at the junction between the healthy outer epithelial layer and the ‘peeling-off’ terminally differentiated cells (Supp.Fig1), suggesting this process as the mechanism by which the membrane can remove unwanted cells and return to its original thin structure.

**Fig.3.**
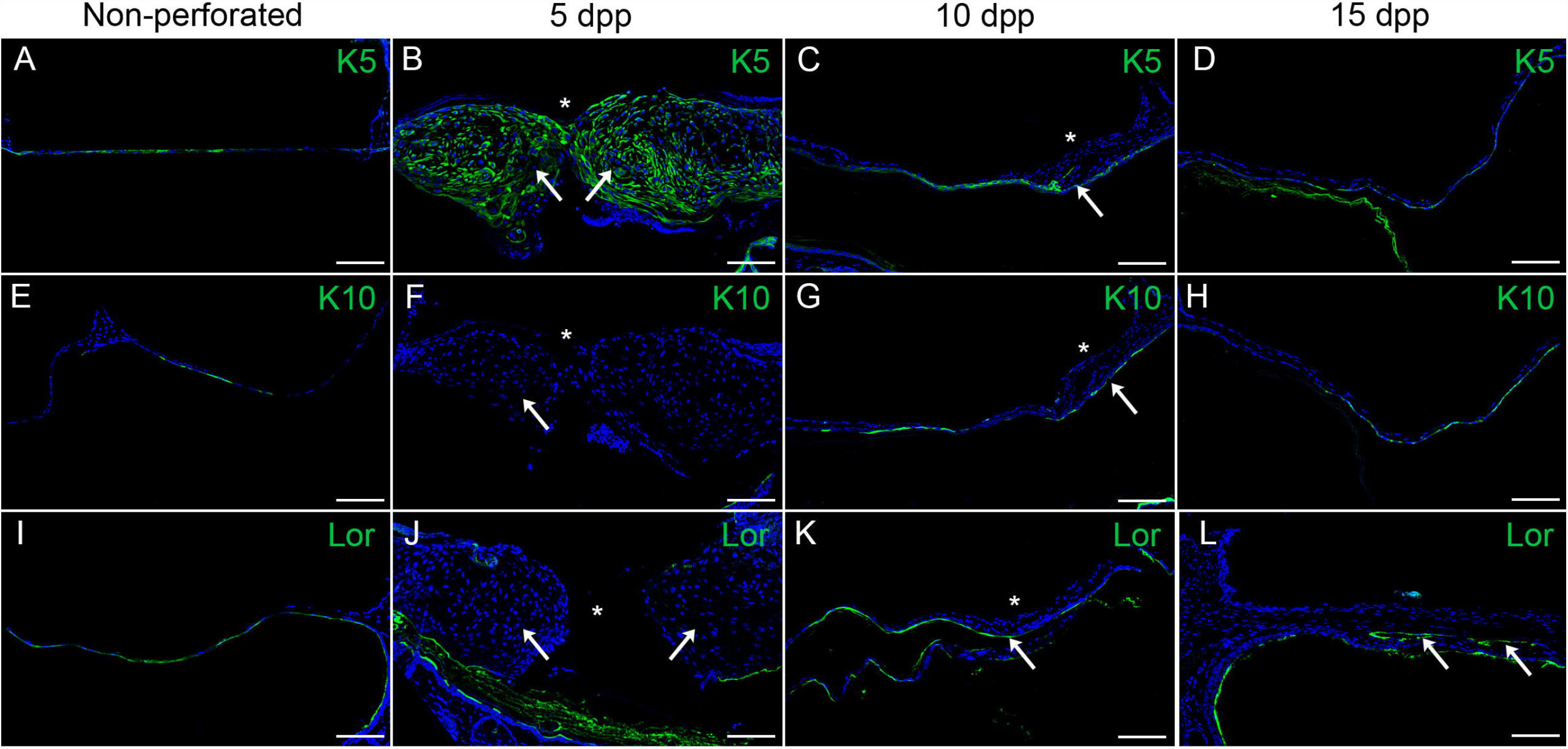
Closure of the perforation by undifferentiated basal cells followed by differentiation. IF for the different epithelial layers markers at the indicated time points post-perforation. A-D) Basal cells marker Keratin 5(K5). E-H) Spinous layer marker Keratin 10 (K10). I-L) Cornified layer marker loricrin. A,E,I) Non-perforated tympanic membranes. B,F,J) 5 days post-perforation. C,G,K) 10 days post-perforation. D,H,L) 15 days post-perforation. Scale Bar: 80μm

### The mesenchymal layers use the epithelial scaffold to cover the wound in tandem with invasion of the vasculature

To identify the response of the middle layer during healing, Wnt1cre;tomato mouse were investigated (Fig.4A-H). During homeostasis, neural crest-derived cells were located around the annulus and the manubrium, with some sparse and isolated mesenchymal cells in the PT (Fig.4A, E, arrows). The sparse nature of the neural crest cells within the main body of the TM, highlights how spaced out the cells are in this thin layer, which is dominated by extracellular matrix (see Fig. 2F). At 5dpp, an increase of tomato+ve cells, suggestive of a thickening of the middle layer, was evident around the manubrium and annulus on the side of the perforation (Fig4.B). At this stage, DAPI positive cells could be observed covering the original perforation site, but no tomato+ve neural crest lineage cells were observed immediately around the closed wound (Fig.4F asterix). By 10dpp (Fig.4C, G) tomato positive cells had migrated into the site of the healing perforation, with a further increase in number or tomato+ve cells by 15dpp (Fig.4D, H). Interestingly, in the unperforated TM, the mesenchymal cells had a clear star shaped morphology with long projections (arrow Fig.4 E arrows, box), while at 15dpp the mesenchyme cells within the healing wound were more compact and elongated (Fig.4 H box).

The middle layer (LP) also contains the vasculature of the TM (Mozaffari et al., 2020). The timing of the invasion of the vasculature was followed in Mesp1cre;tomato mice, where Mesp1 drives tomato in mesodermal derivatives, including endothelial cells (Asrar & Tucker, 2022). In unperforated TMs, blood vessels were observed focused around the annulus and the manubrium, while the centre of the PT was devoid of them (Fig.4 I). At 5 days post perforation, a large change was observed in the blood vessel network, with upregulation of mesodermally-derived cells around the annulus and manubrium on the side of the perforation, mimicking the changes in the mesenchymal layer (Fig. 4J). The centre of the wound, however, remained devoid of any vasculature until 10 dpp (Fig. 4 J,K). By 15dpp the blood vessels had started to disappear/retract from the neawly healed membrane, with a similar location to control conditions (Fig. 4L). The LP and its vasculature, therefore, invade the wound site after the ectoderm. Finally, the response of the inner endoderm layer was followed using *Sox17icre;tomato* mouse line. This line labels endodermal derivatives but also the vasculature (Engert et al., 2009). In unperforated TMs, endoderm derived cells were seen clearly marking the inner layer (Fig. 4 M). At 5dpp there was no evidence of the endoderm creating an early bridge, as has previously been suggested, and unlike the other layers, only a thin layer of endoderm was evident around the wound (Fig.4 N). By 10dpp, the endoderm was observed as a continuous thin layer overlying the other layers on the inside of the drum (Fig. 4 O), with the thin layer being retained by 15dpp (Fig. 4P). Shedding of the inner layer into the middle ear cavity was not evident, in contrast to the marked shedding of the outer epithelial layer into the ear canal (compare Fig. 40,P and 3K,L).

**Fig.4.**
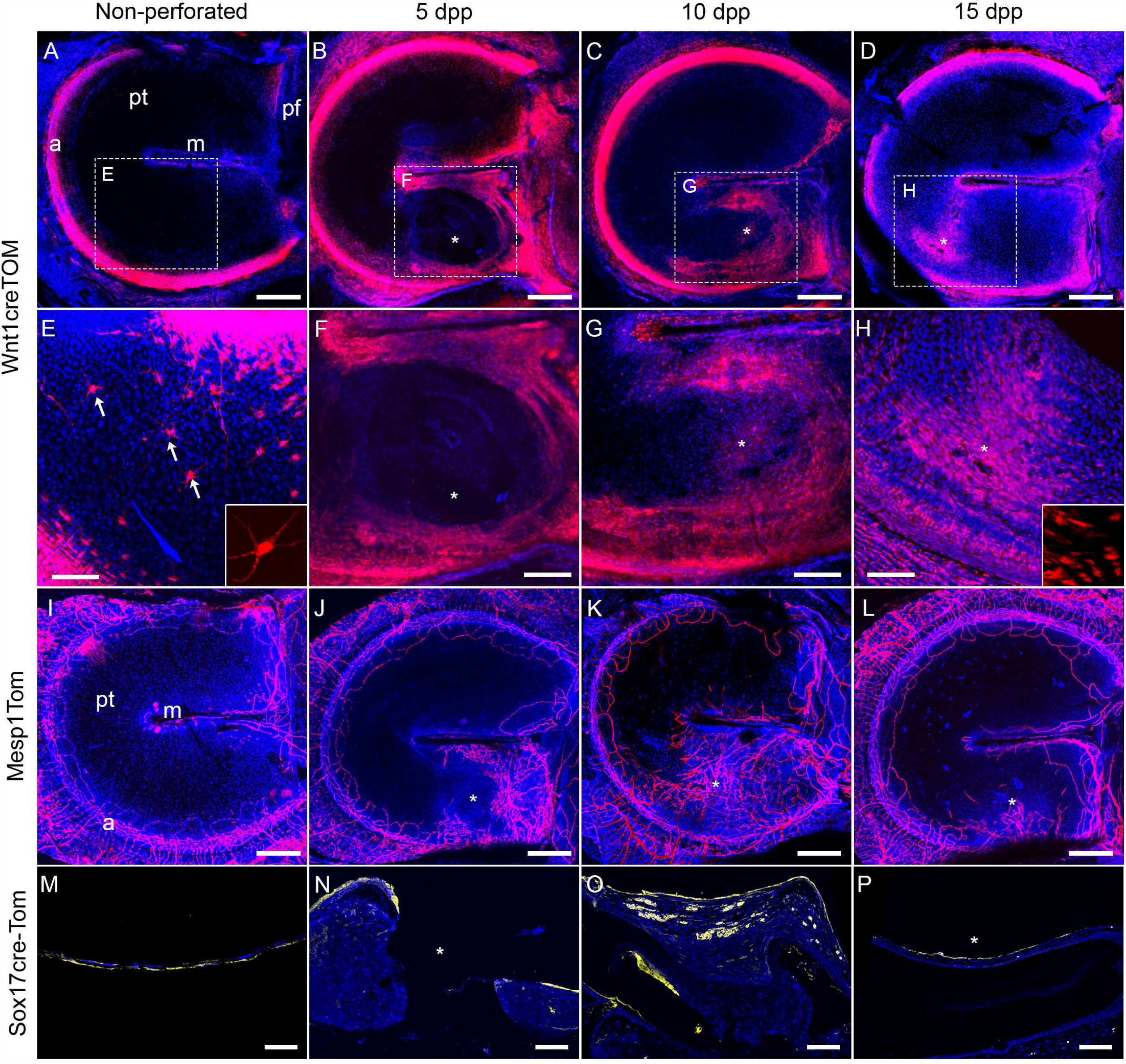
Synchronous healing of perforation by neural crest derived mesenchymal layer, blood vessels and endoderm derived mucosal layer following epithelial scaffold. Whole mount fluorescence of the neural crest derived reporter mouse *Wnt1cre-tdTomato* (A-I) and the mesoderm derived reporter mouse *Mesp1cre-tdTomato* (J-M) in unperforated tympanic membranes (A,F,J) and at 5days (B,G,K), 10 days (C,H,L) and 15 days (E,I,M) post-perforation. Insets in E and H are high magnification of the doted square. M-P) IF for Tomato (in yellow) of transversal sections of the endoderm derived reporter mouse *Sox17icre-tdTomato* at the indicated time points before and after perforation. Nuclei (DAPI) in Blue. pt: pars tensa, pf: pars flacida, m: manubrium, a: annulus. Scale Bar: A-E and J-M, 200 μm. F-I, 100μm. M-P, 80μm. Insets in E,H, 25μm.

### Proliferation of the epithelial layer occurs at specific sites while proliferation of the middle and inner layers occurs across the entire area

To understand the cellular behaviour of the TM during healing, the proliferation pattern of the different layers was analysed. In sections of perforated *Wnt1cre;tom* TMs, PCK co-staining confirmed that the initial bridge was created/being created by an ectodermal spur prior to migration of the middle layer (labelled with tomato) and the mucosal layer (seen here with DAPI) on the opposite site of the PCK positive layer (Fig. 5A). By following the proliferative cells using PCNA staining at 5 dpp, it was observed that the ectodermal proliferation occurred at the annular and manubrial sides of the perforation (Fig.5E, arrow), rather than at the ‘leading edge’, which remained free from proliferation (Fig. 5E). In contrast, the mesenchymal cells next to the annulus were non-proliferation at this time point (Fig.5E). The endodermal cells at the junction of the TM and the middle ear mucosa were also highly proliferative at this stage, this region having been previously noted as an area of label retaining cells (Fig. 5E) (Tucker et al., 2018).

**Fig.5.**
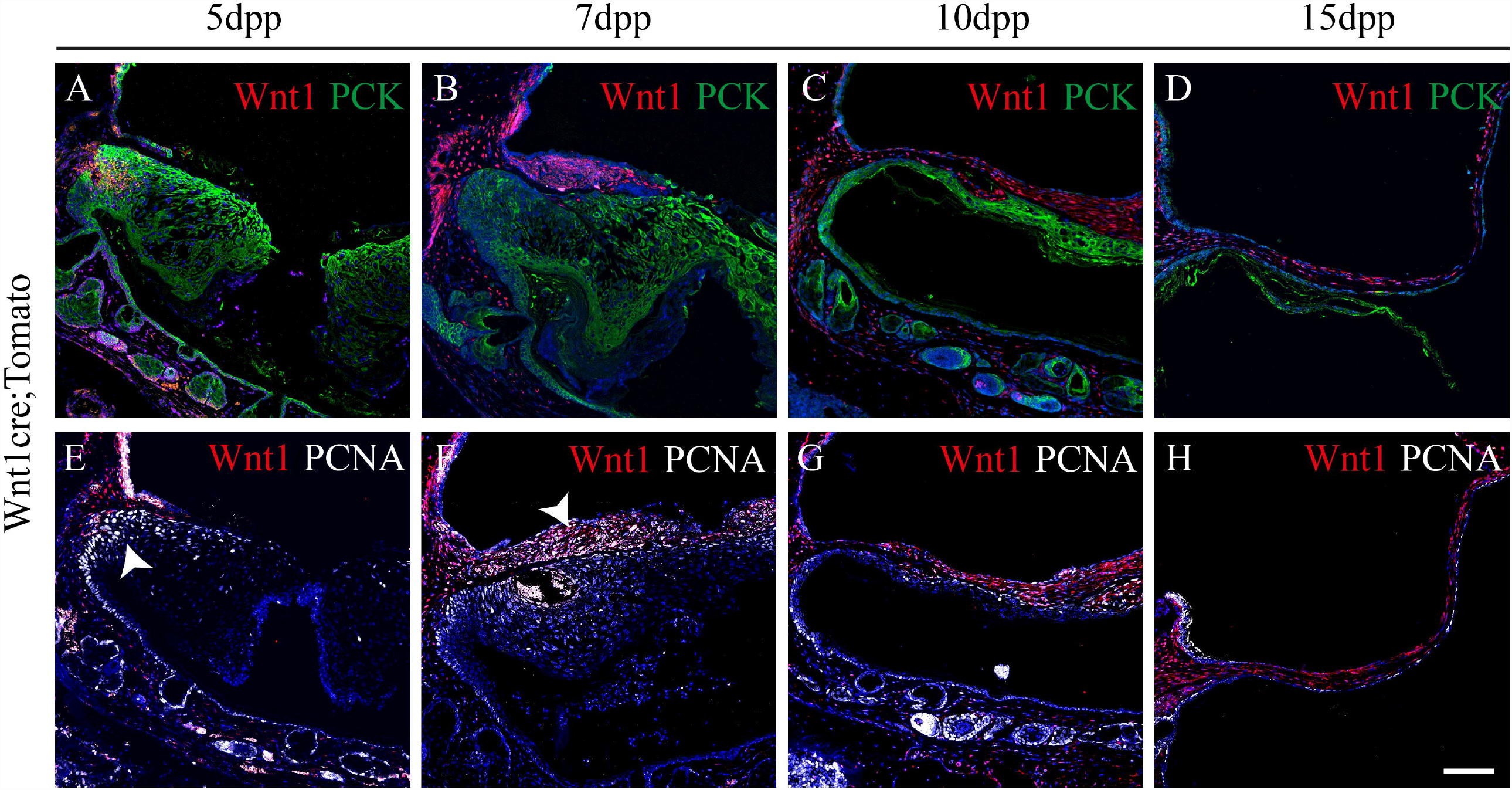
Differential proliferation patterns between layers during healing. A-D) IF for tomato (red) and the epithelial marker pancytokeratin (green) of transversal sections of tympanic membranes at 5 days (A,E), 7 days (B,F), 10 days (C,G) and 15 days (D,H) post perforation (dpp) in the neural crest derived reporter mouse *Wnt1cre-tdTomato*. E-H) IF for RFP (Red fluorescent protein) and the proliferation marker PCNA (white) in consecutive sections as A-D. Nuclei (DAPI) in blue. Arrows indicate proliferating regions. Scale Bar in H = 100μm. Same scale in all other images.

At 7dpp, a thicker mesenchymal layer was evident moving over the epithelial scaffold to reach the wound (Fig.5B). The mesenchyme was highly proliferative at this stage as shown by tomato and PCNA co-staining (Fig.5F), but unlike the epithelial layer, the entire mesenchyme proliferated, rather than this process occurring only near the margins. At this stage, the mucosal layer appeared to move with the underlying neural crest. Proliferative cells were observed along the endoderm (Fig. 5F), despite this layer retaining its single cell structure throughout the healing process (Fig. 5B-D).

By 10dpp, the mesenchymal layer had traversed the entire perforation and continued to proliferate (Fig.5C,G), while proliferation of the ectoderm layer was largely restricted to the basal layer (Fig. 5G). By 15dpp proliferation level had decreased, with ectodermal proliferation restricted to basal cells, and very limited proliferation evident in the mesenchyme or endodermal layers of the TM (Fig. 5D,H).

## Discussion

In this study the contribution of the distinct layers of the TM were assessed during healing. Using immunofluorescence and genetic reporter mouse lines, rather than relying on histological approaches, it has been possible to characterise the contribution of the layers of the TM accurately and reproducibly. After perforation, the outer epithelium was shown to retract by 2 dpp, with the LP and endodermal layers remaining. The outer epithelial layer proliferated at the junction of the annulus and manubrium areas, which pushed the cells over the perforation to form a scaffold by day 5-7. Variation in timing of closure of the gap may depend on subtle differences in position of the perforation or in initial size of the perforation, although variation was reduced for both by visualising the perforation site using an endoscope during surgery. After expansion of the outer epithelial layer, the mesenchymal and inner layers then utilised this scaffold to cover the wound, concomitant with invasion of the vasculature. This was followed by a remodelling step, whereby the outer epithelial scab was lost, and the tissue thinned down to its original, functional size. A schematic highlighting the mechanisms of action of healing is shown in Figure 6.

**Figure 6:**
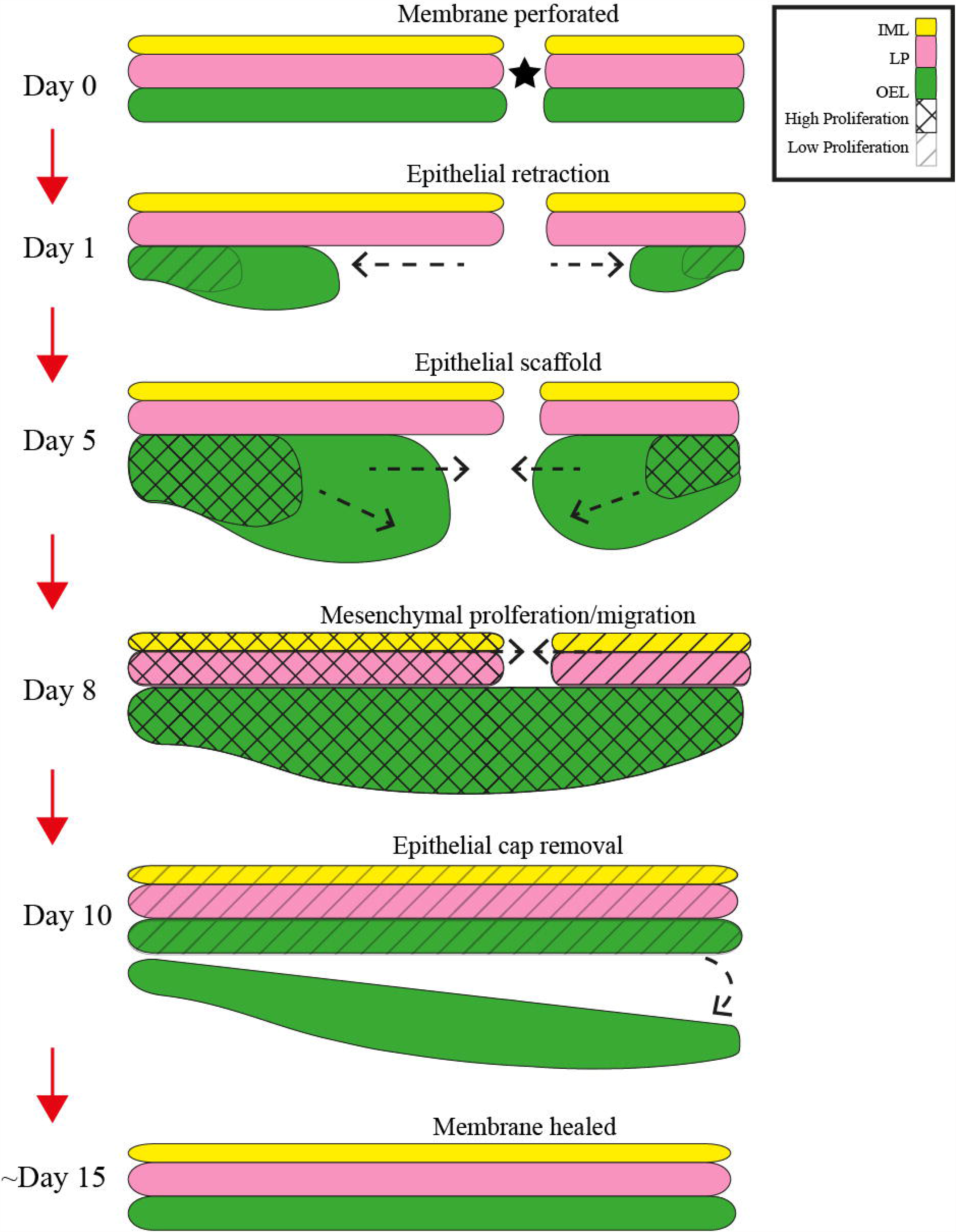
Schematic of the acute healing process of the tympanic membrane. Schematic showing the stages of acute TMP healing. IML (inner mucosal layer) yellow, LP (lamina propria) pink, OEL (outer epithelial layer) green. Shaded regions denote regions of proliferation.

The results highlight that after perforation, a significant retraction of only the epithelial outer layer takes place. This retraction was not immediate, with no evidence at 3 hours post perforation, suggesting it did not involve a reflex contraction, however, it is likely that the retraction was triggered by changes in tension in the usually tight membrane. While there are examples of epithelial retraction involved in healing (e.g., the cornea (Onochie et al., 2019)), it has not been described at the scale observed here. Whether retraction of the epithelium is an integral part of the healing process is unclear, but it did only affect the outer epithelial layer, which is the first tissue to respond to injury, with localized proliferation to push cells over the hole. The outer epithelial tissue had heightened expression of phalloidin compared to the non-retracted tissues, suggesting that actin may help the non-proliferating epithelial cells at the retraction edge to contract over the wound using a purse-string mechanisms.

While previous studies narrowed down areas of proliferation to the side of the TM where the perforation has been made (Chari et al., 2019), we further show that the proliferative zones were restricted (during epithelial scaffold formation) to the annulus and manubrium either side of the perforation with newborn cells pushed away from these regions towards the wound site. The bridging epithelial cells were initially K5+ve, agreeing with previous reports (Chari et al., 2019), and later differentiated into K10+ve and loricrin+ve cells, showing their ability to mature in a matter of days. The ability of the enlarged epithelial layer to ‘thin down’ was explained by the peeling of the loricrin positive cells in layers into the ear canal, from 10dpp to the end of the remodeling phase. During this process the cells to be shed appeared to flatten and with loss of the nucleus, characteristic of cornification, a non-apoptotic form of programmed cell death (Eckhart et al., 2013).

The mesenchymal reaction began after or towards the end of the epithelial scaffold formation at day ∼5. In contrast to the epithelial layer, there were proliferative cells throughout the re-forming mesenchyme from day 7, suggesting that the tissue does not have a ‘pushing mechanism’ as such, and this may be because, unlike the epithelia, the mesenchymal and inner layers can use the epithelial scaffold to travel across the air-borne middle ear. The mucosal layer contributed to the wound closure in tandem with the mesenchymal layer. Interestingly, in contrast to the outer ectodermal layer, the endodermal layer did not increase in thickness at any time point analysed, despite evidence of proliferation. Shedding of tissue during remodelling of the scar, therefore, only occurred on the external ear side of the membrane. In our model, soon after 10dpp the middle layer started to thin down, indicating that the remodelling phase had begun (Santa Maria et al., 2011). How the mesenchymal layer ‘thins’ after its proliferative event is of interest and worth further investigation. As the mesenchymal layer is sandwiched between the outer and inner layers, cells do not have the freedom to ‘peel-off’ into the environment. It is possible that apoptosis has a role, although apoptotic bodies were not evident from DAPI staining (data not shown), or phagocytosis by macrophages, which play important roles to eliminate diseased and damaged tissue in other systems (Hirayama et al., 2017). Vascularisation is important to bring in macrophages and other immune cells, while providing a supply of nutrients during later stages of repair. The increase in the vasculature network occurred in concomitance with the closing middle layer, with a reduction after 10dpp and complete disappearance in the wound site once fully healed. Whether the vessels migrated out or were removed by apoptosis during the remodelling phase is an interesting avenue to follow.

Overall, this research is important as tympanic membrane perforation, and moreover, chronic tympanic membrane perforation has a very high incidence worldwide. Current treatment options for chronic perforations are limited, and surgery focussed. Although, all perforations healed in our mouse model, our research suggests several mechanisms that could disrupt the healing process. The annulus and manubrial regions were heavily involved in the repair process, providing a source of cells for closure, agreeing with these regions containing putative stem cell populations (Tucker et al., 2018). Perforations within these regions, in contrast to the main body of the membrane, would therefore be predicted to heal less well if the stem cell compartment was damaged. In human patients, bilateral repair of holes has been documented (Yamashita, 1985). This could be explained by defects in migration of the neural crest derived layer, which moves in after the outer epithelium. Such migration defects could be a direct issue with the neural crest or could be caused by the epithelial layers over proliferating and physically obstructing movement of the other layers. Finally, it would be interesting to see how infection, such as otitis media (OM), affects the different layers during healing of perforations, which should be possible by crossing the reporter mice to mouse models of OM (Fons et al., 2022). By understanding the contribution of each cell layer in the healing process it may be possible to generate targeted therapeutics for chronic perforations in the future.

## Author contributions

AT conceived the idea and was awarded funding. OD and JF performed the perforations, histology and immunofluorescence and imaged the samples. OD wrote the initial draft of the manuscript, which was edited by JF with final version written by AST. All authors contributed to the article and approved the submitted version.

## Funding

This work was supported by the Medical Research Council (MRC), project grant awarded to AST (MR/R023719/1). OD was funded by the King’s College London MRC doctoral training programme.

## Conflict of interest statement

The authors declare no conflict of interest.

## Data sharing

The data that support the findings of this study are shown here and are available from the corresponding author upon request.

## Acknowledgements

Mesp1cre mice were provided by the RIKEN BRC through the National Bio-Resource project of MEXT, Japan. Sox17-2A-iCre mice were provided by Heiko Lickert.

## Notes

### Competing Interest Statement

The authors have declared no competing interest.

